# The relationship between species richness and ecosystem variability is shaped by the mechanism of coexistence

**DOI:** 10.1101/098384

**Authors:** Andrew T. Tredennick, Peter B. Adler, Frederick R. Adler

## Abstract

Theory relating species richness to ecosystem variability typically ignores the potential for environmental variability to promote species coexistence. Failure to account for fluctuation-dependent coexistence mechanisms may explain observed deviations from the expected negative diversity–ecosystem variability relationship, and limits our ability to predict the consequences of future increases in environmental variability. We use a consumer-resource model to explore how coexistence via the temporal storage effect and relative nonlinearity affects ecosystem variability. We show that a positive, rather than negative, diversity–ecosystem variability relationship is possible when ecosystem function is sampled across a natural gradient in environmental variability and diversity. We also show how fluctuation-dependent coexistence can buffer ecosystem functioning against increasing environmental variability by promoting species richness and portfolio effects. Our work provides a general explanation for variation in observed diversity–ecosystem variability relationships and highlights the importance of conserving regional species pools to help buffer ecosystems against predicted increases in environmental variability.

## INTRODUCTION

MacArthur (1955), Elton (1958), and even Darwin (Turnbull et al. 2013) recognized the potential for compensatory dynamics among species to stabilize ecosystem functioning in fluctuating environments. This idea underlies the “insurance hypothesis” (Yachi & Loreau 1999), which states that ecosystem variability, defined as the coefficient of variation of ecosystem biomass over time, should decrease with diversity because species respond dissimilarly to environmental variation, broadening the range of conditions under which the community maintains function (Loreau 2010). A variety of theoretical models all predict a negative relationship between species richness and ecosystem variability (Lehman & Tilman 2000; Ives & Hughes 2002; Loreau & de Mazancourt 2013), and experimental tests tend to support such a prediction (Tilman et al. 2006; Hector et al. 2010).

However, the ability of biodiversity–ecosystem functioning (BEF) experiments to accurately represent real-world dynamics is debated (Eisenhauer et al. 2016; Wardle 2016). Much of the debate centers on the fact that BEF experimental protocols do not allow species additions from the regional pool to offset species losses in local communities. Theoretical work on diversity–ecosystem variability relationships typically suffers from the same limitation: it recognizes the role of environmental variability in driving population fluctuations which destabilize ecosystems, but ignores the potential for environmental variability to promote species richness and thereby help stabilize ecosystems (Loreau 2010, but see Chesson et al. 2001).

Fluctuating environmental conditions are an important ingredient for stable species coexistence, both in theoretical models (Chesson 2000a; Chesson et al. 2004) and in natural communities (Cáceres 1997; Descamps-Julien & Gonzalez 2005; Adler et al. 2006; Angert et al. 2009). Such “fluctuation-dependent” coexistence emerges most easily when species have unique environmental responses and environmental conditions vary so that each species experiences both favorable and unfavorable conditions, preventing competitive exclusion (Chesson 2000a). Chesson (2000) described the two fluctuation-dependent mechanisms: the storage effect and relative nonlinearity. Both mechanisms operate when environmental variation favors different species at different times. Under the storage effect, this happens because species are competing for resources at different times (and escaping competition in unfavorable periods). Under relative nonlinearity, all species are competing for resources at the same time, but each species alters resource availability in a way that favors its competitors. We describe these mechanisms in more detail below (see **Materials and Methods: Consumer-resource model**).

When coexistence is maintained by a fluctuation-dependent mechanism, an increase in environmental variability might lead to an increase in species richness and, consequently, a decrease in ecosystem variability. However, increasing environmental variability may also increase ecosystem variability by increasing the fluctuations of individual species, regardless of species richness. These countervailing effects of environmental variability present an interesting paradox: while we should expect an increase in environmental fluctuations to increase ecosystem variability, this increase might be buffered if fluctuation-dependent coexistence adds new species to the community. Such a paradox complicates predictions about how ecosystems will respond to predicted departures from historical ranges of environmental variability.

The opposing effects of environmental variability on ecosystem variability might explain the mixed results from observational studies on the diversity–ecosystem variability relationship. Observational tests of the diversity–ecosystem variability relationship, which require sampling across natural diversity gradients, have yielded negative (Hautier et al. 2014), neutral (Valone & Hoffman 2003; Cusson et al. 2015), and positive (Sasaki & Lauenroth 2011) relationships. In a meta-analysis of diversity–ecosystem variability relationships, Jiang & Pu (2009) found no significant evidence for an effect of species richness on ecosystem variability when restricting data to observational studies in terrestrial ecosystems, perhaps because environmental variability varies across natural diversity gradients, affecting both richness and ecosystem variability. The idiosyncratic results of these observational studies contrast with the consistent conclusions from experimental and theoretical work that ignore, or control, the feedbacks between variability and richness.

The gap between theoretical expectations and empirical results of diversity–ecosystem variability relationships might reflect the divergence of theory developed to explain species coexistence and theory developed to explain diversity and ecosystem variability. In his thorough review of the topic, Loreau (2010) cautions that “one of the pieces of the stability jigsaw [puzzle] that is still missing here is the interconnection between community stability and the maintenance of species diversity due to temporal environmental variability.” One reason these two disciplines have diverged is because they have focused on different questions. Diversity–ecosystem variability studies typically ask how ecosystem variability responds to different levels of species richness at a given level of environmental variability (reviewed in Kinzig et al. 2001; Loreau 2010), whereas coexistence studies ask how species richness responds to different levels of environmental variability (Chesson & Warner 1981).

To reconcile these two perspectives, we extend theory on the relationship between species richness and ecosystem variability to cases in which species coexistence explicitly depends on environmental fluctuations and species-specific responses to environmental conditions. We focus on communities where coexistence is maintained by either the temporal storage effect or relative nonlinearity, but not both, using a general consumer-resource model. We use the model to investigate two questions:

1. Does the diversity–ecosystem variability relationship remain negative when species coexistence is maintained by the temporal storage effect or relative nonlinearity?
2. How does increasing environmental variability impact ecosystem variability when coexistence depends on the storage effect or relative nonlinearity?

## MATERIALS AND METHODS

### Consumer-resource model

We developed a semi-discrete consumer-resource model that allows multiple species to coexist on one resource by either the storage effect or relative nonlinearity. In our model, the consumer can be in one of two-states: a dormant state *D* and a live state *N*. The dormant state could represent, for example, the seed bank of an annual plant or root biomass of a perennial plant. Transitions between *N* and *D* occur at discrete intervals between growing seasons, with continuous-time consumer-resource dynamics between the discrete transitions. Thus, our model is formulated as “pulsed differential equations” (Pachepsky et al. 2008; Mailleret & Lemesle 2009; Mordecai et al. 2016). We refer to τ as growing seasons and each growing season is composed of *T* daily time steps, indexed by *t* (*t* = 1,2,3,…, *T*). For example, the notation τ(*t*) reads as: “day *t* within growing season τ.”

At the beginning of growing season τ a season-specific fraction (γ_i_,_τ_) of dormant biomass is activated as living biomass such that

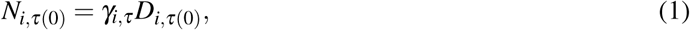
where *i* indexes each species and τ(0) denotes the beginning of growing season τ. Live biomass at the start of the growing season (*N*_*i*,τ(0)_) then serves as the initial conditions for continuous-time consumer-resource dynamics that are modeled as two differential equations:

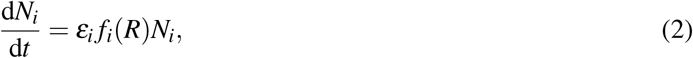

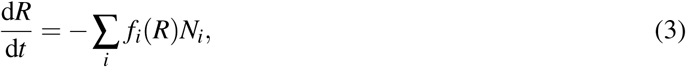
where the subscript *i* denotes species, *N_i_* is living biomass, and ε*_i_* is species-specific resource-to-biomass conversion efficiency. The growth rate of living biomass is a resource-dependent Hill function, 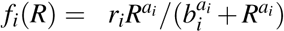, where *r* is a species’ intrinsic growth rate and *a* and *b* define the curvature and scale of the function, respectively. Resource depletion is equal to the sum of consumption by all species.

At the end of the growing season (*t* = *T*), a fraction (α_i_) of live biomass is stored as dormant biomass and a fraction of dormant biomass survives (1 − *η_i_*) to the next growing season, giving the following equation:

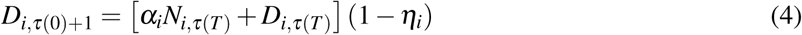
where τ(*T*) denotes the end of growing season τ. We assume remaining live biomass (N*_i_*_,τ(_*_T_*_)_,(1 − α*_i_*)) dies (i.e., this is not a closed system where all biomass must be in either *N* or *D* states). We do not include extinction thresholds, or any other form of demographic stochasticity, under the assumption that we are working with abundant species with generous seed dispersal.

We assume the resource pool is not replenished within a growing season. Resource replenishment occurs between growing seasons, and the resource pool (*R*) at the start of the growing season is *R*_τ(0)_ = *R*^+^, where *R*^+^ is a random resource pulse drawn from a lognormal distribution with mean *μ* (*R*^+^) and standard deviation σ (*R*^+^). Taken all together, we can combine equations 1 and 4 to define the discrete transitions between live and dormant biomass at the end of a growing season. Thus, the initial conditions for each state (*D, N, R*) at the beginning of growing season τ + 1 are:

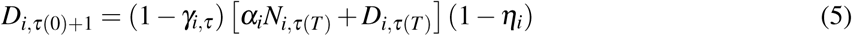

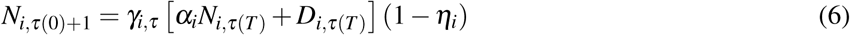

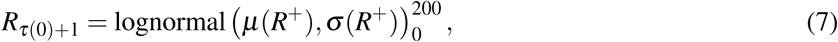
where, as above, τ(*T*) denotes the end of growing season τ and τ(0) + 1 denotes the beginning of growing season τ + 1. The subscript (0) and superscript (200) indicates a lognormal distribution truncated at those values to avoid extreme resource pulses that cause computational problems. We used the function `urlnorm` from the `Runuran` package (Leydold & Hörmann 2015) to generate values from the truncated lognormal distribution. Model parameters and notation are described in Table 1.

**Table 1.**
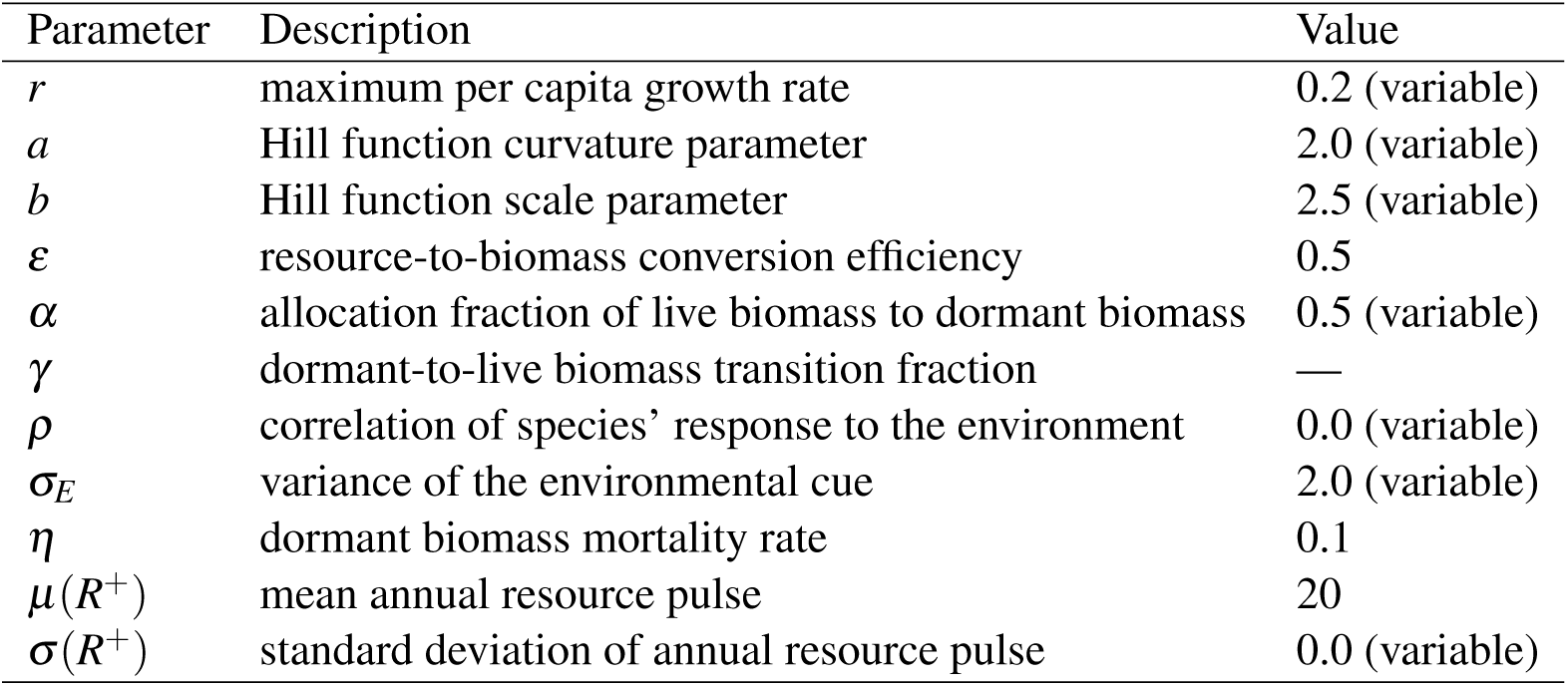
Default values of model parameters and their descriptions. Parameters that vary depending on the mode and strength of species coexistence or depending on species competitive hierarchies are labeled as “variable” in parentheses. The dormant-to-live biomass transition fraction (*γ*) is a function of other parameters, so has no default value.

| Parameter | Description | Value |
| --- | --- | --- |
| $r$ | maximum per capita growth rate | 0.2 (variable) |
| $a$ | Hill function curvature parameter | 2.0 (variable) |
| $b$ | Hill function scale parameter | 2.5 (variable) |
| $\varepsilon$ | resource-to-biomass conversion efficiency | 0.5 |
| $\alpha$ | allocation fraction of live biomass to dormant biomass | 0.5 (variable) |
| $\gamma$ | dormant-to-live biomass transition fraction | — |
| $\rho$ | correlation of species’ response to the environment | 0.0 (variable) |
| $\sigma_E$ | variance of the environmental cue | 2.0 (variable) |
| $\eta$ | dormant biomass mortality rate | 0.1 |
| $\mu(R^+)$ | mean annual resource pulse | 20 |
| $\sigma(R^+)$ | standard deviation of annual resource pulse | 0.0 (variable) |

Our model does not include demographic stochasticity, which can lead to stochastic extinction for small populations as environmental variability increases (Boyce 1992). Previous work has shown how demographic stochasticity and coexistence mechanisms can interact to create a weak “humped-shape” relationship between coexistence time and environmental variability (Adler & Drake 2008), because environmental variability increases coexistence strength and the probability of stochastic extinction simultaneously. We do not consider this potential effect here because our focus is on large populations that would most influence ecosystem functioning.

We limit our analysis to four-species communities because it is exceedingly difficult to get more than four species to coexist via relative nonlinearity without introducing some form of fluctuation-independent mechanism like resource partitioning (Yuan & Chesson 2015), which would then cloud our inference on the effect of fluctuation-dependent coexistence. For consistency, we also constrain our focus to four species communities under the storage effect, but our conclusions apply to more species-rich communities (see Supporting Information section SI.2).

### Implementing the Storage Effect

For the storage effect to operate, we need species-specific responses to environmental variability, density-dependent covariance between environmental conditions and competition (*EC* covariance), and subadditive population growth (Chesson 1994, 2000b). If these conditions are present, all species can increase when rare and coexistence is stable. In the storage effect, rare species increase by escaping the effects of *EC* covariance. Common species will experience greater than average competition (*C*) in environment (*E*) years that are good for them because common species cannot avoid intraspecific competition. However, a rare species can escape intraspecific competition and has the potential to increase rapidly in a year when the environment is good for them but bad for the common species. *EC* covariance emerges in our model because dormant-to-live transition rates (*γ*) are species-specific and vary through time. In a high *γ* year for a common species, resource uptake will be above average because combined population size will be above average. In a year when γ is high for rare species and low for common species, resource uptake will be below average because combined population size will be below average. Subadditive population growth buffers populations against large population decreases in unfavorable years. It is included in our model through a dormant stage with very low death rates, which limits large population declines in bad *E* years.

We generated sequences of (un)correlated dormant-to-live state transition rates (*γ*) for each species by drawing from multivariate normal distributions with mean 0 and a variance-covariance matrix (Σ(*γ*)) of

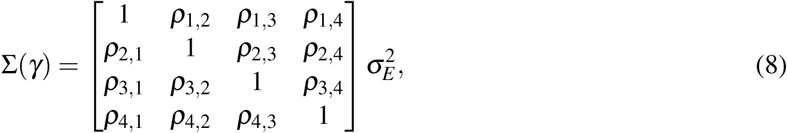
where 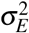 is the variance of the environmental cue and *ρ_i,j_* is the correlation between species *i*’s and species *j*’s transition rates. *ρ* must be less than 1 for stable coexistence, and in all simulations we constrained all *ρ_i,j_*’s to be equal. In a two-species community, the inferior competitor has the greatest potential to persist when *ρ* = −1 (perfectly uncorrelated transition rates). However, in a four-species community the minimum possible correlation among species is -1/3 given our constraints that all *ρ*’s are equal and that Σ(*γ*) must be positive-definite. We used the *R* function mvrnorm to generate sequences of (un)correlated variates E that we converted to germination rates in the 0-1 range: *γ* = *e^E^/* (1 + e*^E^*).

### Implementing Relative Nonlinearity

In the absence of environmental fluctuations, the outcome of competition between two species limited by the same resource is determined by the shape of their resource uptake curves. That is, at constant resource supply, whichever species has the lowest resource requirement at equilibrium (*R*\*) will exclude all other species (Tilman 1982). Resource fluctuations create opportunities for species coexistence because the resource level will sometimes exceed the *R*^*^ of the superior competitor. If the resource uptake curves of each species are relatively nonlinear, then some species will be able to take advantage of resource levels that other species cannot (Chesson 1994).

For example, in Fig. 1C we show uptake curves of two species with different degrees of nonlinearity. Species B has the lowest *R*^*^ and would competitively exclude species A in the absence of environmental fluctuations. But fluctuating resource supplies can benefit species A because it can take advantage of relatively high resource levels due its higher saturation point. Stable coexistence is only possible, however, if when each species is dominant it improves conditions for its competitor. This occurs in our model because when a resource conservative species (e.g., species B in Fig. 1C) is abundant, it will draw resources down slowly after a pulse, and its competitor can take advantage of that period of high resource availability. Likewise, when a resource acquisitive species (e.g., species A in Fig. 1C) is abundant, after a pulse it quickly draws down resources to levels that favor resource conservative species. Such reciprocity helps each species to increase when rare, stabilizing coexistence (Armstrong & McGehee 1980; Chesson 2000a; Chesson et al. 2004).

**Figure 1.**
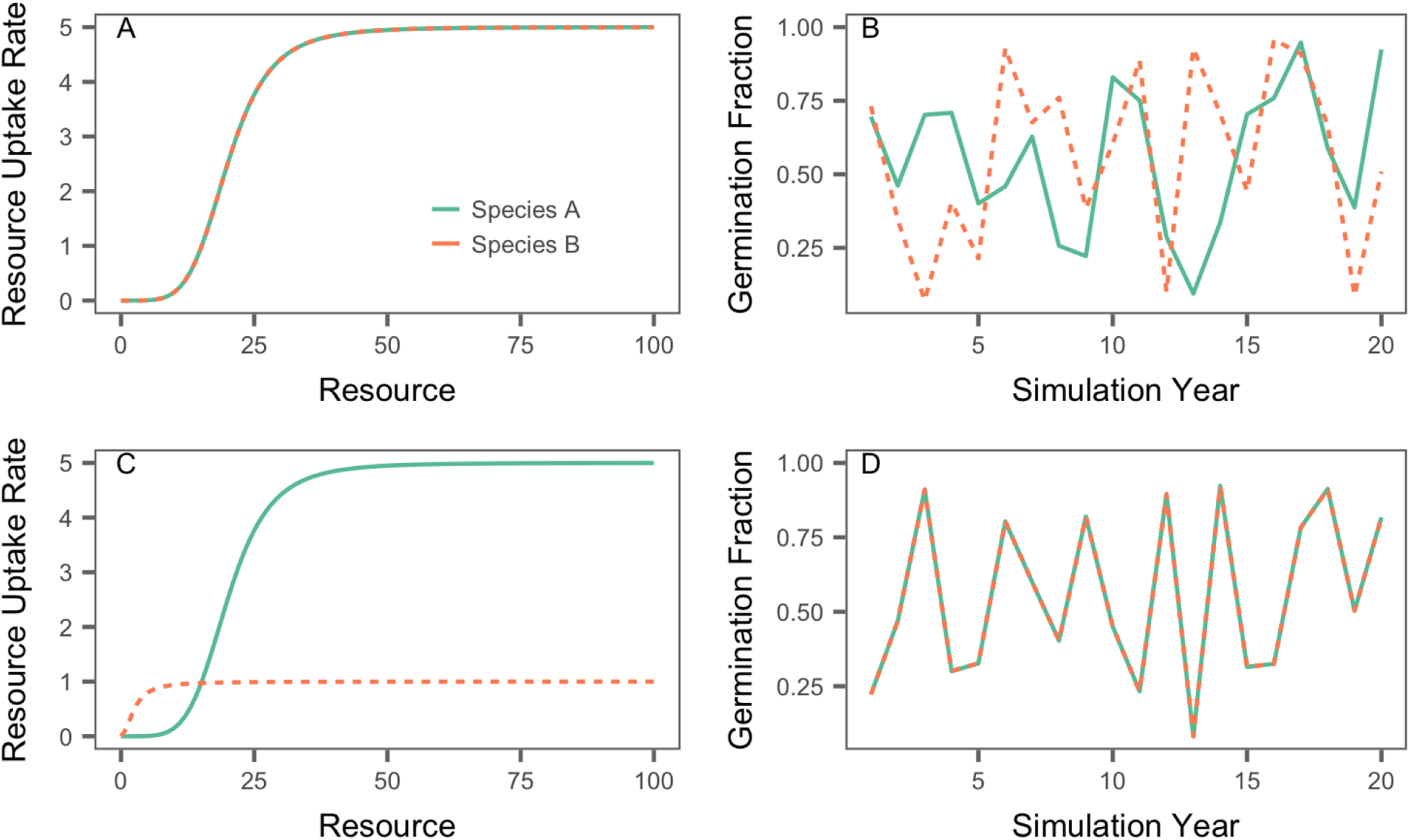
Resource uptake functions and example time series of (un)correlated germination fractions for the storage effect (A,B) and relative nonlinearity (C,D) formulations of the consumer-resource model. The resource uptake functions for both species are equivalent for the storage effect, but their dormant-to-live transition fractions (*γ*) are uncorrelated in time. The opposite is true for relative nonlinearity: the two species have unique resource uptake functions, but their dormant-to-live transition fractions (*γ*) are perfectly correlated in time.

### Numerical simulations

To explore how fluctuation-dependent coexistence can affect the diversity–ecosystem variability relationship, we simulated the model with four species under two scenarios for each coexistence mechanism. First, we allowed the variance of the environment to determine how many species can coexist, akin to a community assembly experiment with a species pool of four species. To do this, we simulated communities with all species initially present across a gradient of annual resource variability for relative nonlinearity (50 evenly-spaced values of σ*_R_* in the range [0, 1.2]) or environmental cue variability for the storage effect (100 evenly-spaced values of 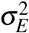 in the range [0, 3]). Second, we chose parameter values that allowed coexistence of all four species and then performed species removals at a single level of environmental variability, akin to a biodiversity–ecosystem function experiment. The two simulation experiments correspond to (i) sampling ecosystem function across a natural gradient of species richness and (ii) sampling ecosystem function across diversity treatments within a site. We refer to the former as a "regional" relationship, and the latter as a “local” relationship. But note that we do not attribute any particular area size to “region”, it is simply any area over which a gradient of environmental variability can emerge.

To understand how increasing environmental variability will impact ecosystem variability when coexistence is fluctuation-dependent, we simulated the model over a range of species pool sizes and environmental cue or resource variability. For each size of species pool (1, 2, 3, or 4 species), we simulated the model at 15 evenly-spaced levels of environmental cue (range = [0.1, 2]) for the storage effect and 25 evenly-spaced levels of resource variability (range = [0, 1.2]) for relative nonlinearity. We also explored the influence of asymmetries in species’ competitive abilities and correlations in species’ environmental responses within the storage effect model. We created competitive hierarchies by making the live-to-dormant biomass fractions (αs) unequal among species. A small difference among values of *α* were needed to create competitive hierarchies because we chose a relatively constrained gradient of environmental cue variance. Larger differences among values of *α* expand the region of coexistence farther along a gradient environmental cue variance. These parameter value choices do not affect our conclusions, but would change the quantitative results.

Under relative nonlinearity, species’ resource response curves (Fig. SI-5) reflect traits that determine the temporal variability of each species’ population growth. “Stable” species achieve maximum resource uptake at low resource levels, but their maximum uptake rates are modest. For these species, population responses to resource fluctuations are buffered. “Unstable” species have very high maximum uptake rates, which they only achieve when resource availability is high, leading to large population fluctuations. The difference in the intrinsic stability of these two kinds of species makes our simulations sensitive to initial conditions. Therefore, we ran two sets of simulations for relative nonlinearity: beginning with either stable or unstable species as a reference point. For example, if species A is the most stable species and species D is the least stable, we ran simulations where A then B then C then D were added to the initial pool of species. We then ran simulations with that order reversed.

All simulations were run for 5,000 growing seasons of 100 days each. We averaged biomass over the growing season, and yearly values of live-state biomass were used to calculate total community biomass in each year. After discarding an initial 500 seasons to reduce transient effects on our results, we calculated the coefficient of variation (CV) of summed species biomass through time, which represents ecosystem variability, the inverse of ecosystem stability. We calculated realized species richness as the number of species whose average biomass was greater than 1 over the course of the simulation. In some cases, realized species richness is less than number of species initialized for a simulation because of competitive exclusion.

For parameters that we did not vary, we chose values that would allow coexistence of all four species at some point along the environmental variability gradients we simulated. Our focus is specifically on communities where fluctuation-dependent coexistence is operating, and making parameters increasingly asymmetric among species typically reduced coexistence strength or made coexistence impossible (Supporting Information section SI.3). Changes in the absolute values of parameters also altered the strength of coexistence, but in no case did altering parameter values change the qualitative results and conclusions we present below (see Supporting Information section SI.3). Parameter values for specific results are given in figure captions. Within-season dynamics were solved given initial conditions using the package deSolve (Soetaert et al. 2010) in R (Team 2013). R code for our model function is in the Supporting Information section SI.1. All model code has been deposited on Figshare (*link*) and is available on GitHub at http://github.com/atredennick/Coexistence-Stability.

## RESULTS

When we allowed the variance of the environment to determine which of four initial species coexisted, similar to a study across a natural diversity gradient, we found a positive relationship between richness and ecosystem variability, defined as the temporal *CV* of total community biomass (Fig. 2A,C). This was true for the storage effect, where coexistence is maintained by fluctuating dormant-to-live transition rates (*γ*), and for relative nonlinearity, where coexistence is maintained by annual resource pulses. The relationship is driven by the fact that increasing environmental variability increases the strength of both coexistence mechanisms (Fig. SI-6). More variable conditions promoted species richness, creating a positive relationship between diversity and ecosystem variability.

**Figure 2.**
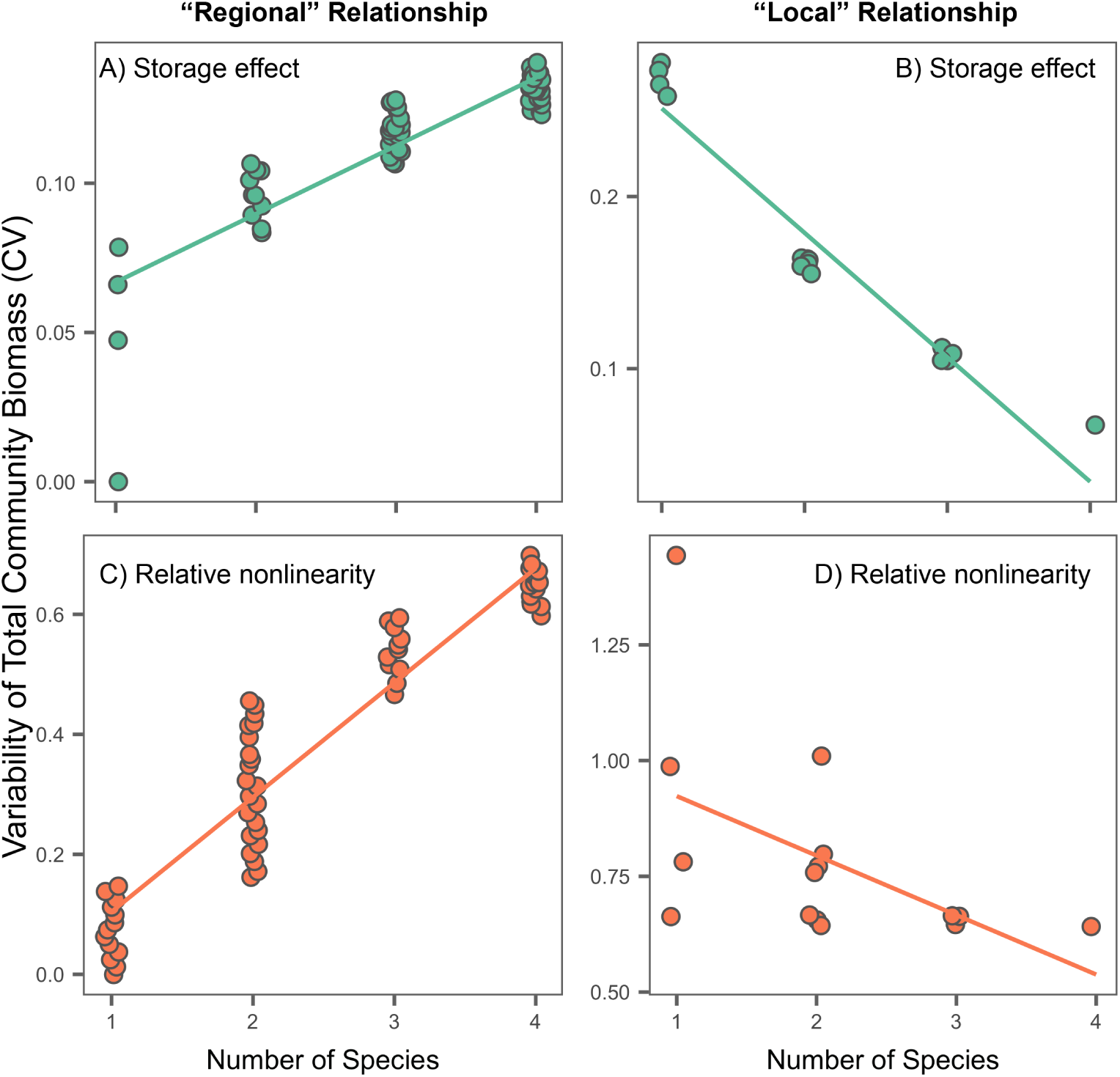
Variability of total community biomass as a function of species richness when coexistence is maintained by the storage effect (A,B) or relative nonlinearity (C,D). Left panels show results from simulations where environmental or resource variance determine the number species that coexist in a community. Right panels show results from simulations where environmental or resource variance is fixed at a level that allows coexistence of all four species, but species are removed to manipulate diversity. The left-hand panels represent “regional” diversity–ecosystem variability relationships across natural diversity gradients, whereas the right-hand panels represent “local” diversity–ecosystem variability relationships. Note that we do not attribute any particular area size to “region”, it is simply any area over which a gradient of environmental variability can emerge. Points are jittered within discrete richness values for visual clarity. Parameter values, where species are denoted by numeric subscripts: (A) *r*_1_ = *r*_2_ = *r*_3_ = *r*_4_ = 0.2, *a*_1_ = *a*_2_ = *a*_3_ = *a*_4_ = 2, *b*_1_ = *b****_2_***= *b*_3_ = *b*_4_ = 2.5, *α*_1_ = 0.5, *α*_2_ = 0.49, *α*_3_ = 0.48, *α*_4_ = 0.47, *ρ*_1_ = *ρ*_2_ = *ρ*_3_ = *ρ*_4_ = 0, σ*_E_* = variable; (B) *r*_1_ = *r*_2_ = *r*_3_ = *r*_4_ = 0.2, *a*_1_ = *a*_2_ = *a*_3_ = *a*_4_ = 2, *b*_1_ = *b*_2_ = *b*_3_ = *b*_4_ = 2.5, *α*_1_ = 0.5, *α*_2_ = 0.49, *α*_3_ = 0.48, *α*_4_ = 0.47, *ρ*_1_ = *ρ*_2_ = *ρ_3_*= *ρ_4_*= -1/3, *Oe* = 4; (C) *r*_1_ = 0.2, *r*_2_ = 1, *r*_3_ = 2, *r*_4_ = 5, *a*_1_ = 2, *a*_2_ = 5, *a*_3_ = 10, *a*_4_ = 25, *b*_1_ = 2.5,*b*_2_ = 20, *b*_3_ = 30, *b*_4_ = 45, *α*_1_ = *α*_2_ = α_3_ = *α*_4_ = 0.5, *ρ*_1_ = *ρ*_2_ = *ρ*_3_ = *ρ*_4_ = 1, σ(*R*^+^) = variable; (D) *r*_1_ = 0.2, *r*_2_ = 1, *r*_3_ = 2, *r*_4_ = 5, *a*_1_ = 2, *a*_2_ = 5, *a*_3_ = 10, *a*_4_ = 25, *b*_1_ = 2.5, *b*_2_ = 20, *b*_3_ = 30, *b*_4_ = 45, *α*_1_ = *α*_2_ = *α*_3_ = *ρ*_4_= 0.5, *ρ*_1_ = *ρ*_2_ = *ρ*_3_= *ρ*_4_ = 1, σ(R^+^) = 1.1.

When we performed species removals but held environmental variability at a level that allows coexistence of all four species, similar to a biodiversity–ecosystem functioning experiment, we found a negative diversity–ecosystem variability relationship (Fig. 2B,D). Scatter around the relationship was small for the storage effect because all species have similar temporal variances. Regardless of species identity, the presence of more species always stabilized ecosystem functioning through portfolio effects. In contrast, scatter around the relationship was larger for relative nonlinearity (Fig. 2D) because species with different resource uptake curves had different population variances. Depending on which species were present, two-species communities were sometimes less variable than three-species communities. Furthermore, the slope of the relative nonlinearity diversity–ecosystem variability relationshp in Fig. 2D is sensitive to species’ traits: the difference among species’ resource uptake determines the spread of single-species communities along the y-axis. This means that the relationship can become flat as species become more similar.

For the storage effect, total community *CV* decreased with species richness at a given level of environmental variability because additional species reduced the temporal standard deviation due to portfolio effects (Fig. SI-7). Mean biomass remained the same because all species had the same resource uptake functions, which was necessary to eliminate any potential effects of relative nonlinearity. Portfolio effects under the storage effect remained strong in an eight-species community, where total community *CV* saturated after addition of the fifth species (Fig. SI-1). For relative nonlinearity, total community *CV* decreased with species richness at a given level of environmental variability because additional species increased mean biomass (over-yielding) and, at higher richness (three to four species), reduced the temporal standard deviation (Fig. SI-7). Mean biomass increased because some species had higher growth rates (Fig. SI-5), increasing total biomass.

To understand how much species additions might stabilize ecosystem functioning as environmental variability increases, we simulated our model over a range of environmental variance and species pool sizes. For both coexistence mechanisms, realized species richness increased with environmental variability and, in some cases, increases in richness completely offset the effect of moderate increases in environmental variability on ecosystem variability (Fig. 3 and 4). More species rich communities were less variable on average and, under the storage effect, they increased in ecosystem *CV* at a slower rate than communities with fewer species (e.g., lower slopes in log-log space; Fig. SI-8). The buffering effect of species richness under the storage effect is also evident in Fig. 2A because the relationship between species richness and ecosystem CV begins to saturate. In fact, ecosystem *CV* remains relatively constant past four species when species have independent responses to the environment (*ρ* = 0; Fig. SI-1).

**Figure 3.**
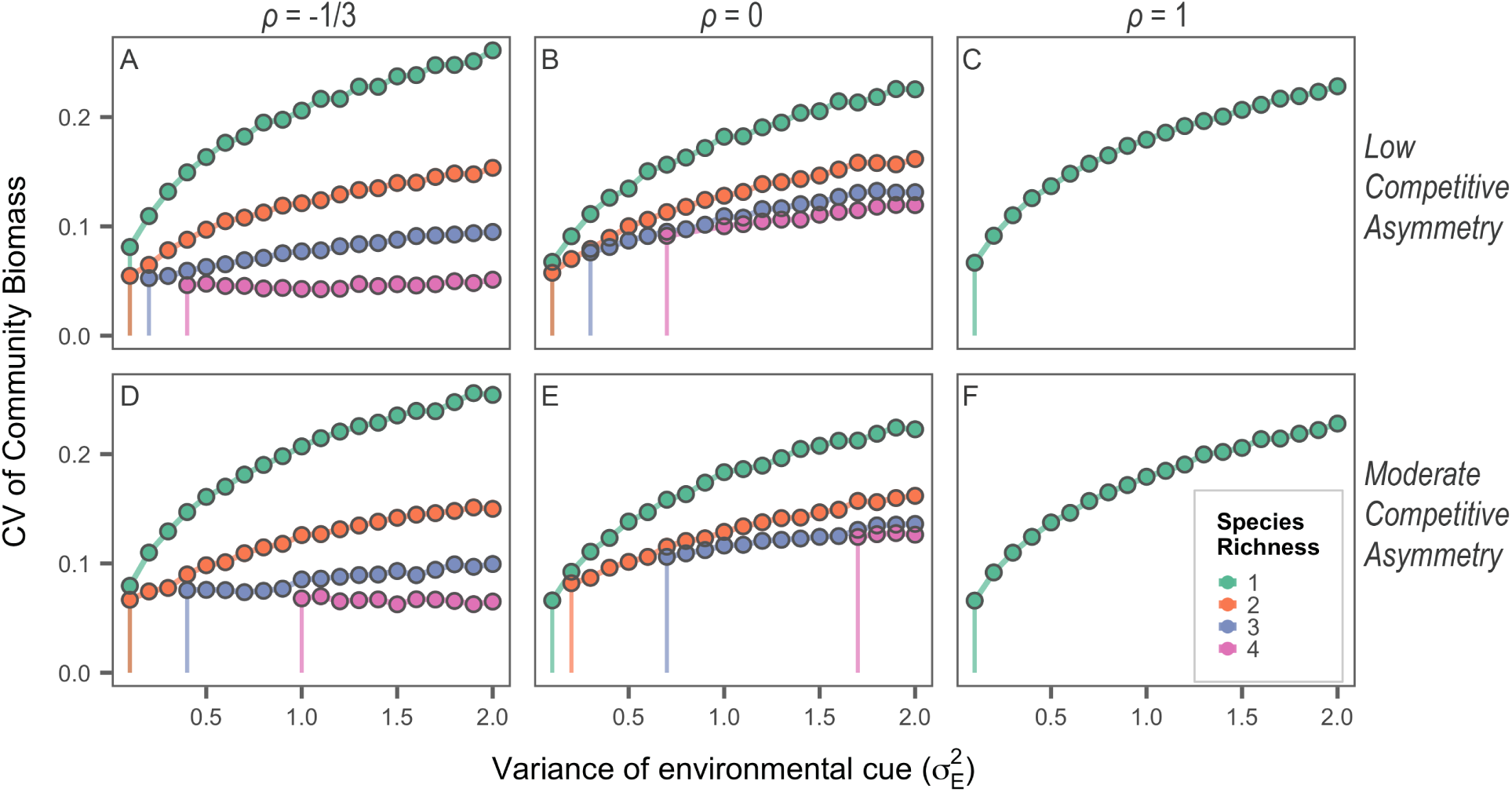
The effect of increasing environmental variability on ecosystem variability when species coexist via the storage effect. Panels (A-C) show simulation results where species have slightly asymmetrical competitive effects, whereas panels (D-F) show results when competition is more asymmetric. Columns show results for different levels of correlations of species’ environmental responses, *ρ*. Colored vertical lines show the magnitude of environmental variability at which each level of species richness first occurs. Parameter values are as in Figure 2A except for as: (A-C) αι = 0.5, α_2_ = 0.495, α_3_ = 0.49, α_4_ = 0.485; (D-F) αι = 0.5, α_2_ = 0.49, α_3_ = 0.48, α_4_ = 0.47.

**Figure 4.**
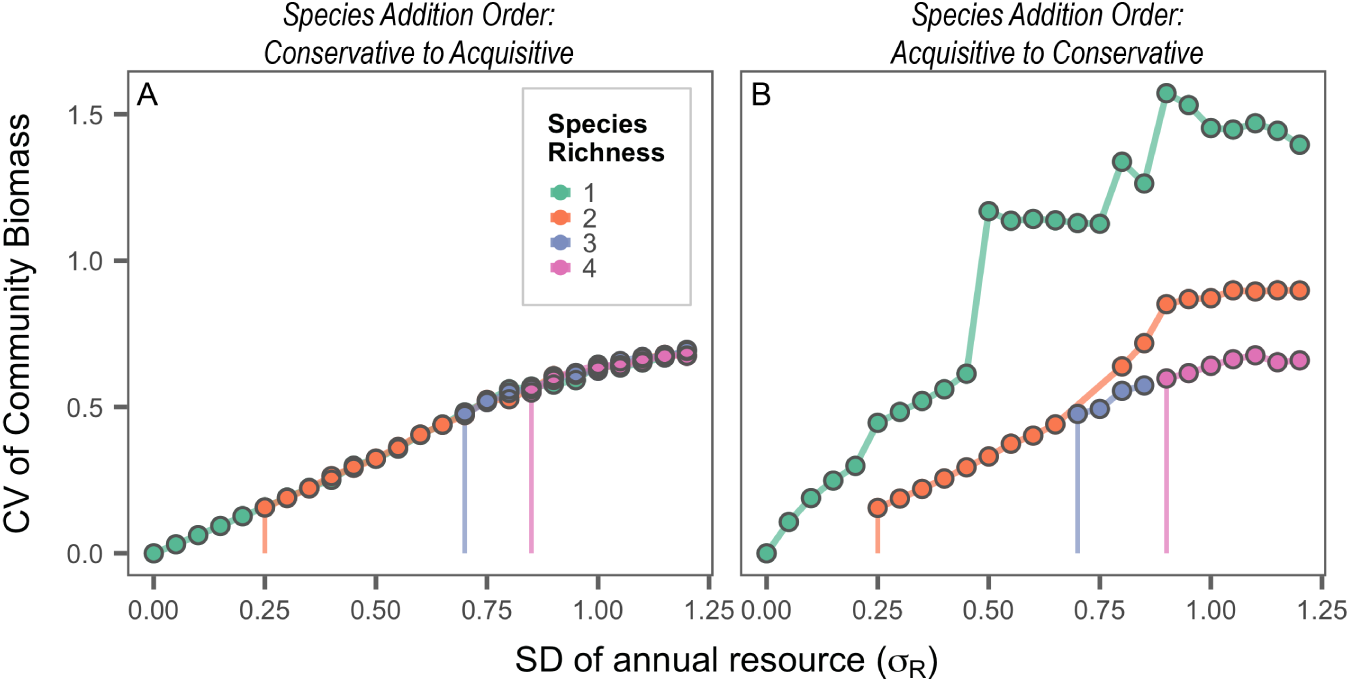
The effect of environmental variability on ecosystem variability when species coexist via relative nonlinearity. (A) The species pool increases from one to four species, with the fourth species being most unstable (e.g., resource conservative to resource acquisitive). Increasing environmental variability (the SD of annual resource availability) allows for greater species richness, but species additions do not modulate the effect of environmental variability on ecosystem variability. (B) The species pool increases from one to four species, with the fourth species being most stable (e.g., resource acquisitive to resource conservative). In this case, increasing environmental variability allows for greater realized species richness and can temper the effect of environmental variability. Parameter values are as in Figure 2C.

The dampening effect of fluctuation-dependent coexistence on increasing environmental variability depends on the specific traits (parameter values) of the species in the regional pool. Under the storage effect, moderately asymmetric competition makes it more difficult for new species to enter the local community, but once they do enter, ecosystem *CV* is similar between communities with low and moderate competitive asymmetries (Fig. 3; compare top and bottom panels). Moderately asymmetric competition does decrease the rate at which ecosystem *CV* increases with environmental variance (Fig. SI-8) because the abundance of inferior competitors is reduced and they do not influence ecosystem *CV* as much as when competitive asymmetry is low. The correlation of species’ environmental responses (*ρ*) also mediates the relationship between environmental variance, species richness, and ecosystem *CV*: lower correlations make it easier for new species to enter the community and contribute to porfolio effects (Fig. 3). When the correlation of species’ environmental responses were as negative as possible (*ρ* = –1/3), ecosystem *CV* of the four-species community was immune to increases in the environmental cue variance (Fig. 3A). However, more extreme increases in the variance of the environmental cue, which increase the number of extremely low or high germination events (i.e., *γ* ≈ 0 or 1; Fig. SI-9), eventually caused ecosystem *CV* to increase in the four species community (Fig. SI-10).

In communities where species coexist via relative nonlinearity, the extent to which species additions buffer ecosystem stability against increases in environmental variability depends on the species traits of immigrating species and the order in which they enter the community. When additional species, which immigrate from the regional pool, are less intrinsically stable than the resident species, ecosystem variability increases at a relatively constant rate even as species are added (Fig. 4A; Fig. SI-11). If more stable species colonize, species additions buffer the ecosystem from increasing environmental variability (Fig. 4B). The “stair-steps” in Fig. 4B emerge because a different species is making up the one-species community at different levels of resource variability, increasing ecosystem CV each time as a more unstable species composes the single species community.

We tested the generality of our results under different parameters by conducting a targeted sensitivity analysis focused on parameter values and asymmetries that most affect species coexistence (Supporting Information section SI.3). Making parameters asymmetric creates competitive hierachies and makes coexistence more difficult. For the storage effect, this is evident in Fig. 3 where we made live-to-dormant transition rates (αs) asymmetric, and the same result occurred when we made dormant mortality rates (*η* s) asymmetric (Fig. SI-2). Furthermore, increasing the absolute value of dormant mortality also made coexistence more difficult, but results were qualitatively similar to those above (Fig. SI-3). For relative nonlinearity, altering the absolute value of the mean resource pulse shifts the region of coexistence across a gradient of resource variability (Fig. SI-4). But, again, qualitatively similar results emerge when species *can* coexist. In general, altering any parameter in isolation will make coexistence easier or harder at any given level of environmental variability. Thus, our results are robust in the sense that the qualitative patterns we present will always emerge for parameter values that permit species coexistence over some range of envrionmental variability. In other words, our results are only sensitive to whether or not fluctuation-dependent coexistence is operating.

## DISCUSSION

Theory developed for biodiversity–ecosystem function experiments emphasizes that increases in species richness should reduce ecosystem variability. Consistent with theoretical expectations from models in which species coexistence is maintained by fluctuation-independent mechanisms and with results from biodiversity–ecosystem functioning experiments, our model of fluctuation-dependent species coexistence (also see Chesson et al. 2001) produced a negative diversity–ecosystem variability relationship (Fig. 2B,D). This agreement is encouraging because empirical evidence for fluctuation-dependent coexistence is accumulating (Pake & Venable 1995; Cáceres 1997; Descamps-Julien & Gonzalez 2005; Adler et al. 2006; Angert et al. 2009; Usinowicz et al. 2012) and species almost certainly coexist by some combination of fluctuation-independent (e.g., resource partitioning) and fluctuation-dependent mechanisms (Ellner et al. 2016). By extending theory to communities where species richness is explicitly maintained by temporal variability, we have gained confidence that experimental findings are generalizable to many communities. In local settings where environmental variability is relatively homogeneous, reductions in the number of species should increase the variability of ecosystem functioning, regardless of how coexistence is maintained.

When we allowed communities to assemble at sites across a gradient of environmental variability, we discovered a positive relationship between species richness and ecosystem variability (Fig. 2A,C). While surprising when viewed through the lens of biodiversity–ecosystem functioning theory and experimental findings, such a relationship is predicted by theory on coexistence in fluctuating environments. Environmental variability is a prerequisite for the storage effect and relative nonlinearity to stabilize coexistence (Chesson 2000a). These mechanisms can translate increased variability into higher species richness (Fig. SI-6), but the increase in environmental variability also increases ecosystem variability. However, the apparent saturation of the relationship in Fig. 3A suggests that the portfolio effects that buffer ecosystems against environmental variability, and inherently emerge under the storage effect, get stronger as more species are able to coexist. Indeed, the relationship between species richness and ecosystem *CV* completely saturates under the storage effect in more species rich communities (Fig. SI-1). This suggests neutral diversity–ecosystem variability relationships are possible due to the storage effect.

Our results may explain why deviations from the negative diversity–ecosystem variability relationship often come from observational studies (Jiang & Pu 2009). Observational studies must rely on natural diversity gradients, which do not control for differences in environmental variability among sites. If species richness depends on environmental variability, it is entirely possible to observe positive diversity–ecosystem variability relationships. For example, DeClerck et al. (2006) found a positive diversity–ecosystem variability when sampling conifer richness and the variability of productivity across a large spatial gradient in the Sierra Nevada, across which environmental variability may have promoted coexistence. Sasaki and Lauenroth (2011) also found a positive relationship between species richness and the temporal variability of plant abundance in a semi-arid grassland. Their data came from a six sites that were 6 km apart. While Sasaki and Lauenroth explained their results in terms of dominant species’ effects (e.g., Thibaut & Connolly 2013), our findings suggest an alternative explanation: each site may have experienced sufficiently different levels of environmental variability to influence species coexistence.

While our modeling results show that fluctuation-dependent coexistence can create positive diversity– ecosystem variability relationships, whether such trends are detected will depend on the particular traits of the species in the community and the relative influence of fluctuation-dependent and fluctuation-independent coexistence mechanisms. Thus, our results may also help explain observational studies where no relationship between diversity and variability is detected. For example, Cusson et al. (2015) found no relationship between species richness and variability of abundances in several marine macro-benthic ecosystems. Many of their focal ecosystems were from highly variable intertidal environments. If coexistence was at least in part determined by environmental fluctuations, then the confounding effect of environmental variability and species richness could offset or overwhlem any effect of species richness on ecosystem variability. This may be particularly common in natural communities, where environmental fluctuations can help promote species coexistence even in cases where fluctuation-independent coexistence mechanisms are most important (Ellner et al. 2016). Previous theoretical work showed how environmental variation can mask the effect of species diversity on ecosystem productivity when sampling across sites (Loreau 1998). Our mechanistic model extends that conclusion to ecosystem variability.

Whether coexistence is fluctuation-independent or fluctuation-dependent becomes especially important when we consider how ecosystem variability responds to increasing environmental variability. In the fluctuation-independent case, species richness is essentially fixed because the niche and fitness differences that determine coexistence are not linked to environmental variability. Therefore, increasing environmental variability will always increase ecosystem variability by increasing the fluctuations of individual species’ abundances. When coexistence is fluctuation-dependent, however, the outcome is less certain. By simulating communities with different species pool sizes across a gradient of environmental variability, we showed that species gains due to increasing environmental variability can buffer the direct effect of environmental variability on ecosystem variability (Figs. 3 and 4).

We relied on numerical simulations of a mechanistic model to reach our conclusions, meaning our results could be sensitive to the specific parameters values we chose. In a targeted sensitivity analysis (Supprting Information section SI.3), we found that our qualitative results are robust so long as specific parameter combinations allow fluctuation-dependent species coexistence (by either the storage effect or relative nonlinearity). Investigating the case in which both the storage effect and relative nonlinearity operate remains a future challenge.

Overall, our results lead to two conclusions. First, when predicting the impacts of increasing environmental variability on ecosystem variability, the mechanism of coexistence matters. Fluctuation-dependent coexistence can buffer ecosystems from increasing environmental variability by promoting increased species richness. Whether our theoretical predictions hold in real communities is unknown and requires empirical tests. Doing so would require manipulating environmental variability in communities where coexistence is known to be fluctuation-dependent, at least in part. Such data do exist (Angert et al. 2009), and a coupled modeling-experimental approach could determine if our predictions hold true in natural communities.

Second, whether local fluctuation-dependent communities can receive the benefit of additional species depends on a diverse regional species pool. If the regional pool is not greater in size than the local species pool, than ecosystem variability will increase with environmental variability in a similar manner as in fluctuation-independent communities because species richness will be fixed (Fig. 5A,B). Metacommunity theory has made clear the importance of rescue effects to avoid species extinctions (Brown & Kodric-Brown 1997; Leibold et al. 2004). Here, instead of local immigration by a resident species working to rescue a species from extinction, immigration to the local community by a new species rescues ecosystem processes from becoming more variable (Fig. 5C,D). Thus, our results reinforce the importance of both local and regional biodiversity conservation. Just as declines in local species richness can destabilize ecosystem functioning (Tilman et al. 2006; Hector et al. 2010; Hautier et al. 2014), species losses at larger spatial scales can also increase ecosystem variability. Wang & Loreau (2014) show that regional ecosystem variability depends on regional biodiversity through its effects on beta diversity and, in turn, the asynchrony of functioning in local communities. Our results show that, when coexistence is fluctuation-dependent, regional biodiversity declines could also affect local ecosystem functioning by limiting local colonization events that could be possible under scenarios of increasing environmental variability (Fig. 5).

**Figure 5.**
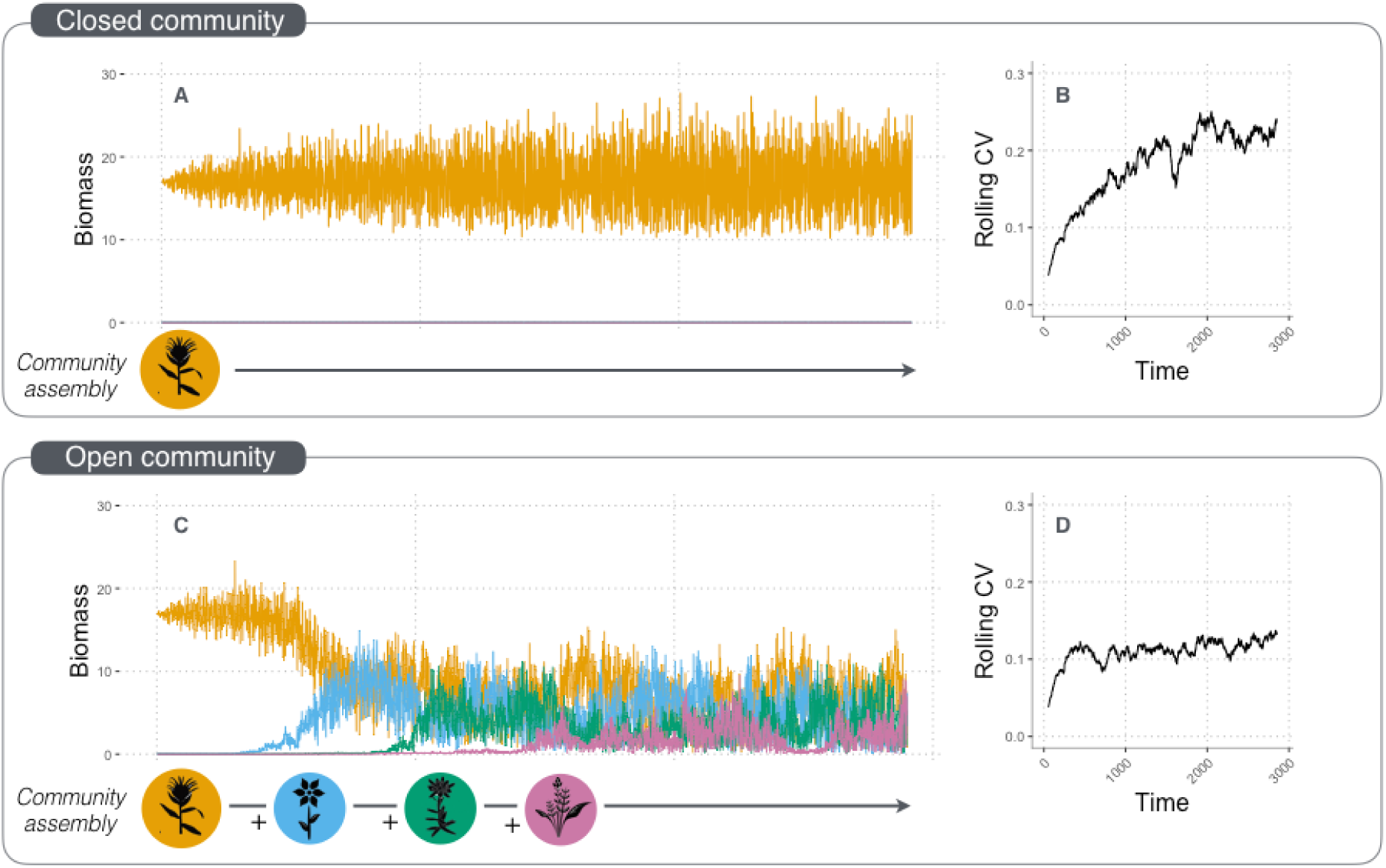
Example of how species additions under increasing environmental variability can buffer ecosystem stability when species coexist via the storage effect. Environmental variability (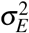) increases linearly with time on the *x*-axis. (A) Time series of species’ biomass (colored lines) in a closed community where colonization of new species is prevented and (B) its associated coefficient of variation (Rolling CV; calculated over 100-yr moving window) through time. (C) Time series of species’ biomass in an open community where colonization by new species from the regional pool of four species becomes possible as environmental variation increases. The trajectory of total biomass CV in the open community (D) asymptotes at lower variability than in the closed community (B) due to the buffering effect of species richness. Parameter values are as in Figure 2A except for *α*s: α_1_ = 0.5, α_2_ = 0.494, α_3_ = 0.49, α_4_ = 0.483.

## ACKNOWLEDGMENTS

The National Science Foundation provided funding for this work through a Postdoctoral Research Fellowship in Biology to ATT (DBI-1400370) and a CAREER award to PBA (DEB-1054040). The support and resources from the Center for High Performance Computing at the University of Utah are gratefully acknowledged. We thank four anonymous reviewers for providing detailed and thoughtful comments that greatly improved the paper.

